# Tanycyte Bmal1 sex-specifically regulates weight gain and hypothalamic neurogenesis in female mice

**DOI:** 10.1101/2025.04.21.649851

**Authors:** Daniel Maxim Iascone, Pavel Pivarshev, Jianing Yang, Mariela Lopez Valencia, Sara B Noya, Hongtong Lin, Ron C Anafi, Joseph L Bedont, Amita Sehgal

## Abstract

The hypothalamic radial-glia-like tanycyte population plays important and intertwined roles in feeding and metabolism, reproduction, and seasonality. Although these processes are circadian-regulated and clock genes reportedly show robust cycling along the 3rd ventricle, the role of the clock in tanycytes has not yet been examined. We report here that clock genes cycle with much higher amplitude in ventral tanycytes compared to more dorsal ependymocytes of the 3rd ventricle, and that specific disruption of the tanycyte clock can be achieved by adult *Bmal1* deletion using the *RaxCreER* driver. Adult tanycyte *Bmal1* deletion did not affect circadian rhythms of wheel-running and sleep, but did inhibit weight gain on high-fat diet in female mice. Altered tanycyte-derived hypothalamic neurogenesis, which can regulate feeding and weight gain by contributing new neurons to nearby feeding-relevant nuclei, is one mechanism that likely contributes to this phenotype. Fate mapping studies showed that female mice have higher baseline tanycyte-derived neurogenesis than males, with many of the resulting neurons localizing to the feeding-relevant arcuate nucleus. Female but not male mice show reduced tanycyte-derived arcuate neurogenesis after adult tanycyte *Bmal1* deletion and an increased percentage of newborn arcuate neurons take on a feeding-suppressing POMC neuropeptidergic fate. Thereby, skewing of feeding and satiety promoting fates link the weight homeostasis and neurogenesis effects. Together, our data establish tanycyte Bmal1 as a sexually dimorphic regulator of weight homeostasis, likely mediated at least in part by a female-specific neurogenesis effect in the feeding circuitry.

## Introduction

The circadian clock organizes the physiology and behavior of most lifeforms on Earth into 24-hour rhythms aligned with the solar cycle. This organization begins in most single cells, where a molecular clock driven by an oscillatory transcription/translation feedback loop regulates tissue-specific clock-controlled gene expression throughout the daily cycle. Core mammalian clock components include the heterodimeric transcription factors Bmal1 and Clock, which cooperatively promote expression of their own negative regulatory *Per* and *Cry* genes, ultimately leading to inhibition of Bmal1:Clock driven transcription (Yi et al., 2022). At the circuit level, light input adjusts circadian phase in cells of the suprachiasmatic nucleus (SCN) in the brain, aligning clock-controlled genes that regulate SCN activity to the solar cycle, which in turn orchestrates systemic autonomic, hormonal, and body temperature rhythms that entrain peripheral clocks to light (Astiz et al., 2019; Hastings et al., 2018). Ultimately, this organizes whole-body circadian rhythms in physiology and behavior, including sleep, feeding, and metabolism.

The health impacts of these rhythms are salient in modern society. Widely available artificial lighting and refrigeration facilitate mis-timed exposure of most humans to circadian entrainment cues like light and food at evolutionarily unprecedented levels (Nordhaus, 1996). Moreover, many people live on “flipped” circadian schedules, with >15% of the US workforce employed at shift work hours outside of a standard diurnal schedule (McMenamin, 2007). Elevated risk of obesity and metabolic disease in shift workers suggests not only direct health impacts from their extreme circadian misalignment (Scheer et al., 2009), but also more subtle contributions of mis-timed circadian inputs to our society-wide epidemic of these conditions. This underlines the importance of understanding how circadian regulation is coupled to metabolism.

Such connections are widespread, with metabolism-relevant genes well represented among the ∼15% of the mouse transcriptome under circadian control across tissues (Guo et al., 2005; Mohawk et al., 2012). One cell population of particular interest in linking circadian rhythms with metabolism is hypothalamic tanycytes. These radial glia-like cells line the floor of the 3rd ventricle (3V) proximal to tuberal hypothalamic nuclei involved in feeding and metabolism: the ventromedial hypothalamus (VMH), arcuate nucleus (ARC) and median eminence (ME). Tanycytes play a number of roles in metabolism that are likely under some level of circadian control, including detection and import of peripheral metabolic hormones like leptin, and gating of the central release of systemic TRH, CRH, and sex hormones (Balland et al., 2014; Friedman, 2019; Müller-Fielitz et al., 2017; Prevot et al., 2018).

The evidence linking tanycytes to circadian clocks has thus far been indirect. The 3V possesses high-amplitude and photoperiod sensitive circadian rhythms (Guilding et al., 2009; Lein et al., 2007; Yasuo et al., 2008), and tanycytes are critical downstream effectors of photoperiodically coded melatonin release, acting to coordinate breeding competence, feeding behavior, and metabolism in a seasonally adaptive manner (Wood and Loudon, 2014). The SCN also signals to tanycytes to gate ARC access to peripheral sugars, mediating circadian modulation of feeding and metabolism (Rodríguez-Cortés et al., 2022). But the tanycyte clock’s role in regulating feeding and metabolism has not previously been directly assessed, in part due to the difficulty of achieving its specific and efficient deletion.

In this manuscript, we focus on the role of tanycytes as a source of new adult-born neurons that modify the feeding circuitry in the ARC and ME, in a diet-dependent manner (Chaker et al., 2016; Haan et al., 2013; Lee et al., 2014, 2012; Surbhi et al., 2021; Yoo and Blackshaw, 2018). Initially inspired by analysis of published tanycyte transcriptomics, here we report the effects of RaxCreER-mediated adult deletion of the essential clock component *Bmal1* in hypothalamic tanycytes We observed a sexually dimorphic reduction in weight gain, specifically reducing weight gain in female mice on a high-fat diet (HFD), without altering rhythms in locomotor activity or sleep. Altered weight homeostasis was instead associated with a female-specific decrease in tanycyte-derived hypothalamic neurogenesis, and increased anorexigenic POMC^+^ fate among remaining adult-born, tanycyte-derived ARC neurons. Together, this suggests that tanycyte Bmal1, likely via its role in the cell-autonomous circadian clock, regulates weight homeostasis by influencing adult tanycyte neurogenesis.

## Methods

### Animal Housing, Care, and Genotyping

Combinations of the following alleles were used for most studies: *RaxCreER* (Pak et al., 2014), *Bmal1* floxed allele (JAX #007668), *Bmal1* constitutive null allele (JAX #009100), and Ai14 fluorescent reporter (JAX #007914). Genetically modified mice were provided by Dr. Seth Blackshaw (*RaxCreER*) and Jackson Laboratories (other mutant lines). All lines not received on a C57BL6/J background were back-crossed at least 5 times to C57BL6/J, and wild-type C57BL6/J mice were used for the *in situ* hybridization (ISH) timecourse. Genotyping was conducted by Transnetyx, Inc using proprietary automated genotyping methods. Except where otherwise noted, mice were on a 12hr:12hr vivarium light:dark cycle.

### *In situ* hybridization: clock gene circadian timecourse

Male mice were entrained to a 12hr:12hr light:dark cycle in custom circadian cabinets (Phenome Technologies), and released into constant darkness for 24hrs before beginning CT4-CT24 tissue collections on the second day in constant darkness, as previously described (Bedont et al., 2018; Shimogori et al., 2010) (Bedont et al., 2018; Shimogori et al., 2010). Briefly, mice were sacrificed under dim red light, and their brains were collected fresh frozen in OCT (Tissue-Tek) and stored at −80C. 25um sections were collected on a Leica CM3050 cryostat, dry mounted on Superfrost Plus slides, and stained by chromogenic *in situ* hybridization with partially hydrolyzed riboprobes. Brain sections containing the tuberal hypothalamus were imaged on a ThermoFisher EVOS M7000 microscope, and ImageJ was used to quantify densitometry in anatomically defined β tanycyte-, α tanycyte-, and ependymocyte-rich portions of the 3rd ventricle. We then subtracted the intensity of nearby signal-poor tissue on each section to control for background color. At circadian expression troughs, low magnitude negative values were occasionally computed. For each probe, a scaling factor sufficient to raise the lowest negative value to 0 was added across the dataset to maintain relative differences among timepoints. 5 brain sections per mouse were quantified and averaged for each data point. Circadian analysis of ISH data was performed using the JTK_CYCLE algorithm within the MetaCycle R package (Hughes et al., 2010; Wu et al., 2016).

Riboprobes were generated from the following constructs:

–*Per2*: Accession #AI838843, PCR amplified with T3+T7 primers, riboprobe run off with T7 polymerase –*Bmal1*: *Bmal1* Exon 8 was PCRed with primers (forward: ATGCAGAACACCAAGGAAGG, reverse: CTTCCTCGGTCACATCCTA), and cloned into pCRII-TOPO. PCR amplification was done with T7+Sp6 primers, and riboprobe was runoff with T7 polymerase.

### Tamoxifen Treatment

To induce Cre-dependent *Bmal1* deletion and tdTomato expression for lineage tracing, *RaxCreER*^*/+*^*;Bmal1*^*lox/null*^*;Ai14*^*/+*^ (Tan^*Bmal1* KO^) experimental and littermate control mice for all experiments were fed on one of the following pre-irradiated and vacuum-packed diets: recombination inducing red-dyed 250mg/kg tamoxifen diet (Inotiv/Envigo TD.130856) or control 2016 diet (Inotiv/Envigo 2916.cs) for 5 weeks, beginning at P30-40. Mice were weighed weekly during this time around mid-afternoon (∼ZT7-10 on vivarium light cycle), and mice dropping more than 20% of their starting body weight were euthanized to prevent suffering. Except where otherwise noted, this was followed for all groups by 4 weeks on control 2016 diet, to allow recovery time for the tamoxifen-fed group before beginning other experimental manipulations.

### High-fat Diet Treatment and MRI

Mice were weighed at the end of their recovery period after tamoxifen or control diet to establish a baseline weight, then moved to new cages with HFD (Inotiv/Envigo TD.06414: 60% calories from fat). Mice were then weighed weekly for 12 weeks, consistently around mid-afternoon (∼ZT7-10 on vivarium light cycle). At the end of this time, mice were transferred to a satellite facility, where they underwent MRI measurement of fat:lean mass body composition on an EchoMRI-500 Body Composition Analyzer early in the morning (∼ZT2-4 on vivarium light cycle). A majority of mice from all experimental groups were then sacrificed at ZT12 for *Per2* ISH analysis.

### Wheel-running Recordings

Wheel running activity was measured as previously described (Bedont et al., 2014). Briefly, mice were transferred to individual cages with a running wheel (Actimetrics PT2-MCR2) and *ad libitum* food and water, and entrained to a strict 12hr:12hr light:dark cycle in circadian cabinets (Actimetrics PT2-CCM1 with added running wheel power rails). After allowing at least a week for photoentrainment, wheel running activity was recorded for at least 10 days in 12:12LD, after which constant darkness was begun at lights off and wheel running activity was recorded for an additional 15 days. Recording was logged and circadian parameters were analyzed using Clocklab recording and analysis software (Actimetrics).

### Piezo Sleep Recordings

Mice were entrained to a strict 12hr:12hr light:dark cycle in circadian cabinets in their home cages for at least a week before beginning the experiment. Mice were then transferred to fresh cages, and placed onto PiezoSleep Mouse Behavioral Tracking System recording platforms (Flores et al., 2007; Mang et al., 2014). Mice were given at least 3 days to adapt, followed by at least 4 days of recording that were averaged for recording sleep/wake behavior.

### Tanycyte Adult Neurogenesis Lineage Tracing

For lineage tracing of tanycyte-derived adult born cells, tanycytes in Tan^*Bmal1* KO^ mice and littermate controls were labeled through tdTomato expression induced by TAM chow administration for 35 days starting at P30-40. Starting at P65-75, mice were placed back on control 2016 chow for an additional 35 days to allow tanycyte-born cells to achieve a mature cell fate. Animals were then anaesthetized with isoflurane before intracardiac perfusion with PBS and 4% PFA (Electron Microscopy Sciences). Brains were fixed in 4% PFA overnight at 4C, and 100μm coronal brain sections sampling the entire hypothalamus were obtained using a vibrating microtome (Leica VT1200S). Brain sections were stained with 1:5000 Hoescht in PBS for 15 minutes to label cell nuclei before being mounted on microscope slides. Hoechst staining patterns within hypothalamic nuclei were then used to identify brain sections corresponding to Bregma −1.35mm, −1.55mm, −1.75mm, and −1.85mm for tanycyte-derived cell counts (Figure 2A). TdTomato^+^ neurons within the arcuate nucleus and all tdTomato^+^ astrocytes within these brain sections were counted for analysis. Astrocytes were distinguished from neurons on the basis of their characteristic “star-like” morphology (Figure 2A). For cell count comparisons between control and Tan^*Bmal1* KO^ mice, female and male mice were each normalized to littermate controls of the same sex.

For neuronal subtype fate mapping experiments, mice were placed on control 2016 chow for 90 days after TAM chow administration to measure the fate acquisition of adult-born feeding neurons that persist throughout the 12-week experimental window in which we investigated weight gain on HFD. PFA-perfused brains were fixed in 4% PFA overnight at 4C and then cryoprotected in 30% sucrose in PBS gently rocking overnight at 4C until they were no longer buoyant. Following cryoprotection, 25 μm coronal brain sections were collected on a Leica CM3050 cryostat, with every 5th section mounted onto alternating Superfrost Plus slides (to create multiple subseries sampling the entire hypothalamus).

### Immunohistochemistry

Fluorescent immunostaining was performed as previously described (Surbhi et al., 2021). In brief, brains were permeabilized with 1X PBS + 0.2% Triton X-100. Antigen retrieval was performed with 10mM sodium citrate buffer, with slides incubated at 95C for 5 minutes. Slides were blocked with 5% bovine serum albumin, 10% normal donkey serum, and 0.2% Triton X-100 in 1X PBS for 2 hours. Primary antibodies, rabbit anti-POMC (1:5000, Phoenix Pharmaceuticals, Cat# H-029-30, RRID:AB_2307442) and goat anti-NPY (1:200, Novus, Cat# NBP1-46535, RRID:AB_10009813) were diluted in blocking solution and slides were incubated overnight at 4C. Sections were washed with 1X PBS and incubated with secondary antibodies Alexa Fluor® 647 Donkey Anti-Rabbit IgG (Abcam) and Alexa Fluor® 488 Donkey Anti-Goat IgG (Abcam) diluted 1:250 in blocking solution for 2 hours at room temperature. Slides were washed with 1X PBS and stained with 1:1000 Hoescht in 1X PBS for 5 minutes to label cell nuclei prior to coverslipping with Prolong Diamond antifade mounting media (Fisher). Sections were imaged on Leica STELLARIS 8 confocal microscope (Leica Microsystems) and all immunofluorescent analyses were performed blinded and using ImageJ (NIH).

### Sequencing Analysis

For our analysis single cell RNA-Seq data from Campbell et al. 2017, we selected cells annotated as “Tanycyte1” and “Tanycyte2” for our tanycyte analysis and cells annotated as “Ependymo” for our ependymocyte analysis. Differential expression analyses were performed between the high fat diet group (HFD) and the normal chow group (Ch10) using the Seurat function FindMarkers with default parameters. We then filtered for significant genes using the thresholds adjusted p-value < 0.2 and absolute log_2 fold change > 0.25. Finally, we performed pathway analyses on the significantly DE genes using the function DEenrichRPlot on databases “GO_Biological_Process_2023” and “GO_Molecular_Function_2023”.

### Statistics

GraphPad Prism 10.0 was used for statistical analyses. For comparisons between two experimental groups, unpaired two-tailed Student’s t tests were used. For comparisons between more than two experimental groups, a two-way ANOVA test with a Holm-Sidak multiple-comparison post hoc analysis was performed to compare the differences between individual groups. For comparisons between more than two experimental groups over time, a three-way ANOVA test was used. A p value of less than 0.05 was considered statistically significant. The statistical tests, n (number of animals), and p values for each dataset are provided in the figure legend that accompanies the data. Experiments were performed and quantified by investigators who were blinded to the genotype and treatment of the mice, and were unblinded once summary data were ready to be prepared.

## Results

### Tanycytes but not ependymocytes show diet-induced expression changes in putative neurogenesis and circadian clock-relevant genes

To gauge the likelihood of the tanycyte clock influencing systemic responses to metabolic challenge, we mined existing transcriptomic datasets (Campbell et al., 2017). In this study, mice were fed either HFD or a low-fat diet for one week, and the transcriptomes of hypothalamic cell types including 3V tanycytes and neighboring ependymocytes were probed. Tanycytes (but not less-specialized ependymocytes) showed robust diet-driven changes in gene expression (Figure 1A-B).

**Figure 1.**
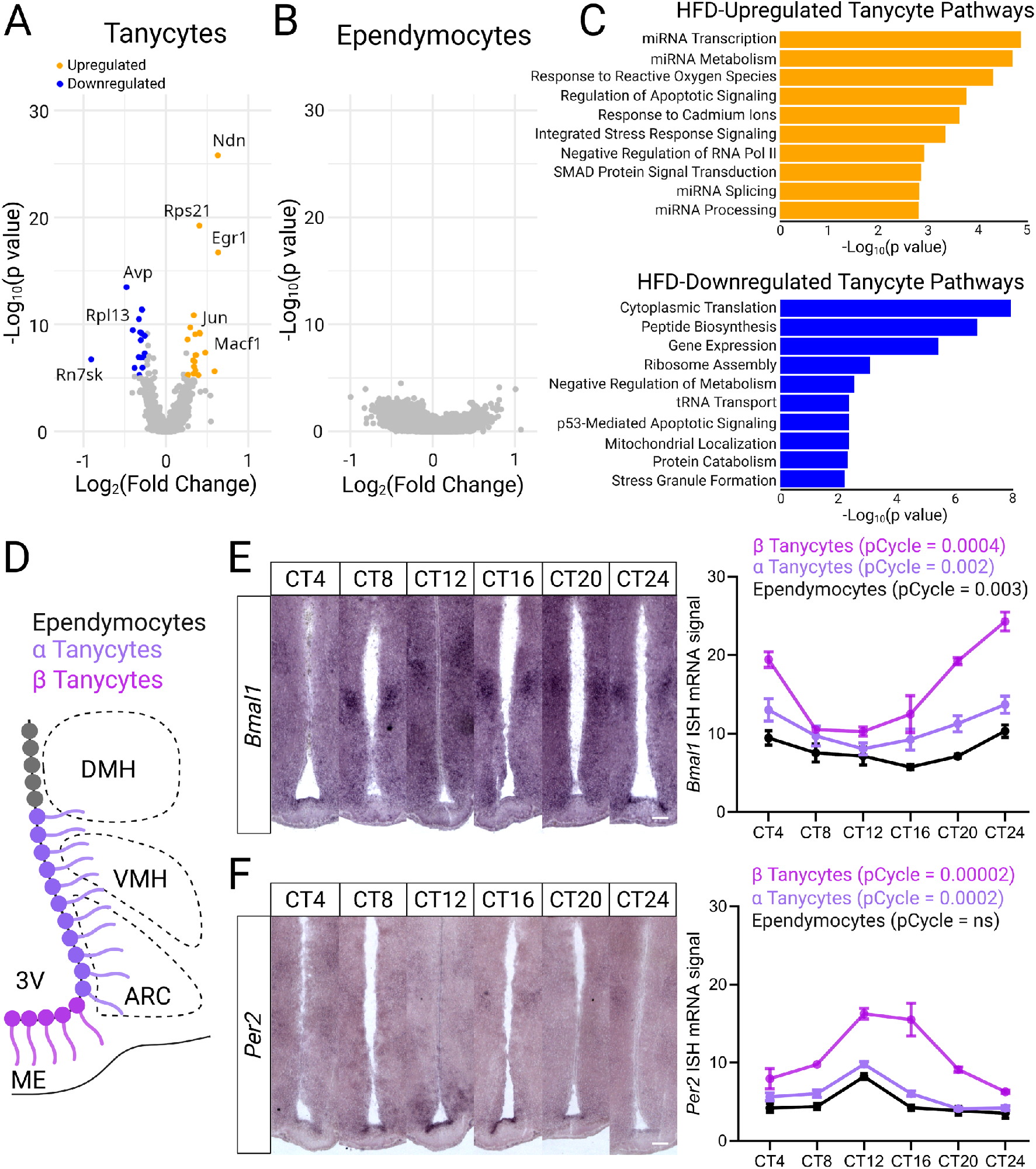
Diet-induced gene expression and circadian oscillation of core clock genes in tanycytes and ependymocytes. (**A-B**) Volcano plots showing differentially expressed genes (DEGs) in tanycytes (A) and ependymocytes (B) from mice fed HFD vs. LFD for 1 week (dataset from Campbell et al. 2017). (**C**) Pathway analysis of significantly diet-regulated tanycyte DEGs from (A) using Gene Ontology enrichment analysis. (**D**) Coronal diagram of mouse hypothalamus indicating cell populations spanning the third ventricle. (**E-F left**) Representative *in situ* hybridization (ISH) images of *Bmal1* (B) and *Per2* (C) mRNA in tanycytes and ependymocytes across circadian time (CT; n = 3-4 mice/CT). Scale bar = 150µm. **(E-F right)** ISH quantification. Mean ± SEM, tested for significant circadian rhythmicity (pCycle) by JTK-cycle, ns = not significant.

Prior literature on highly up-regulated individual transcripts (e.g. *Necdin, Rps21, Egr1*) suggested an inhibitory effect of dietary fat on hypothalamic neurogenesis, consistent with HFD effects on ARC but not ME neurogenesis (Lee et al., 2014, 2012). Both *Necdin* and *Rps21* are anti-proliferative (Török et al., 1999; Yoshikawa, 2021), and Egr1 stimulates IGF receptor signaling, which specifically inhibits tanycyte neurogenesis and self-renewal (Chaker et al., 2016; Ma et al., 2012). Members of the senescence-promoting AP-1 complex including *Jun* and *FosB* were also up-regulated (Martínez-Zamudio et al., 2020). Several transcripts are also associated with aspects of neuronal differentiation, including *Necdin* (differentiation and survival of postmitotic neurons) (Yoshikawa, 2021), *Macf1* (neurite outgrowth and migration) (Moffat et al., 2017), *Jun* (associated with post-mitotic neurons more than neural precursors in adult neurogenesis) (Kawashima et al., 2017), and *Rps21* (most enriched ribosomal factor in dendrites) (Fusco et al., 2021). Consistent with a potential anti-proliferative effect of HFD in tanycytes, Gene Ontology analysis identified changes in cellular pathways associated with ATF4 signaling, including upregulation of reactive oxygen species-associated pathways and the integrated stress response as well as downregulation of cytoplasmic translation and ribosome assembly (Figure 1C) (Frank et al., 2010; Pakos-Zebrucka et al., 2016).

Importantly, our top hits also strongly suggested links to the circadian clock. Egr1 drives liver *Per1* expression and is rhythmically expressed with specific sensitivity to food reward in the brain (Correa-da-Silva et al., 2021; Herichová et al., 2017), while Necdin binds directly to Bmal1 and promotes its stability (Lu et al., 2020). Together, these data suggested a potentially important role for the cell-autonomous circadian clock in modulating tanycyte neurogenic responses to dietary fats, in part by intervening in tanycyte neurogenesis.

### Tanycyte circadian rhythms are ventrally enriched and susceptible to deletion by adult RaxCreER activation

Hypothalamic tanycytes have 4 subpopulations: α1/2 and β1/2. Briefly, β tanycytes line the ME and play prominent roles in blood-brain barrier function, peripheral nutrient sensing, and gating hormone release, while α tanycytes line the ARC and VMH and facilitate cross-talk of these nuclei with the cerebrospinal fluid (Rodriguez et al., 2005). The remaining dorsal extent of 3V is largely composed of non-tanycyte ependymocytes (Figure 1D). While circumstantial literature evidence suggests clock gene enrichment in ventral β and perhaps α2 tanycyte populations, to our knowledge the relative contributions of 3V sub-populations have not previously been directly quantified (Bedont et al., 2020; Guilding et al., 2009; Lein et al., 2007; Yasuo et al., 2008). We tested this by sampling *Bmal1* and *Per2* clock gene transcripts by *in situ* hybridization in anatomically-defined β tanycytes (ME-adjacent), α tanycytes (ARC/VMH-adjacent), and non-tanycyte ependymocytes (remainder of 3V) across circadian time in wild-type mice. As expected, *Bmal1* and *Per2* rhythms were dorsoventrally patterned, with the most robust rhythmicity in β tanycytes (Figure 1D-E).

These data supported targeting tanycyte Bmal1 with RaxCreER^T2^, which is primarily expressed in ARC/ME-adjacent β and α2 tanycytes when activated in adulthood (Pak et al., 2014; Yoo et al., 2020). We tested several tamoxifen induction paradigms targeting the essential clock gene *Bmal1* in *RaxCreER*^*/+*^*;Bmal1*^*lox/lox*^ and *RaxCreER*^*/+*^*;Bmal1*^*lox/-*^ mice (data not shown), and found that a 5-week induction with 250mg/kg tamoxifen (TAM) chow on a *Bmal1*^*lox/null*^ background was needed to consistently blunt the tanycyte cell-autonomous clock in adulthood (Figure S1A). The genetic combination of *RaxCreER* and *Bmal1*^*lox/null*^ will hereafter be referred to as tanycyte *Bmal1* knockout (Tan^*Bmal1* KO^) mice. At least 4 weeks were allowed on control chow after TAM treatment to mitigate direct TAM effects on behavior and physiology.

### Loss of tanycyte *Bmal1* sex-specifically reduces weight gain on high-fat diet (HFD) in female but not male mice

Because tanycyte neurogenesis has consistently been linked to weight gain, feeding, and metabolism (Chaker et al., 2016; Haan et al., 2013; Lee et al., 2014, 2012; Surbhi et al., 2021; Yoo and Blackshaw, 2018),we tested whether tanycyte *Bmal1* knockout affected weight gain on HFD. Control *Bmal1*^*lox/+*^ or experimental Tan^*Bmal1* KO^ mice previously pre-treated with either CreER-activating TAM diet or control 2016 diet were subsequently fed on HFD for 12 weeks. Female mice exhibited main effects of genotype (p<0.0001) and previous TAM/control treatment (p<0.05), with a highly significant interaction (p<0.0001). Indeed, while TAM drove *increased* weight gain in genetic control mice, TAM actually *reduced* weight gain in Tan^*Bmal1* KO^ mice (Figure 2).

**Figure 2.**
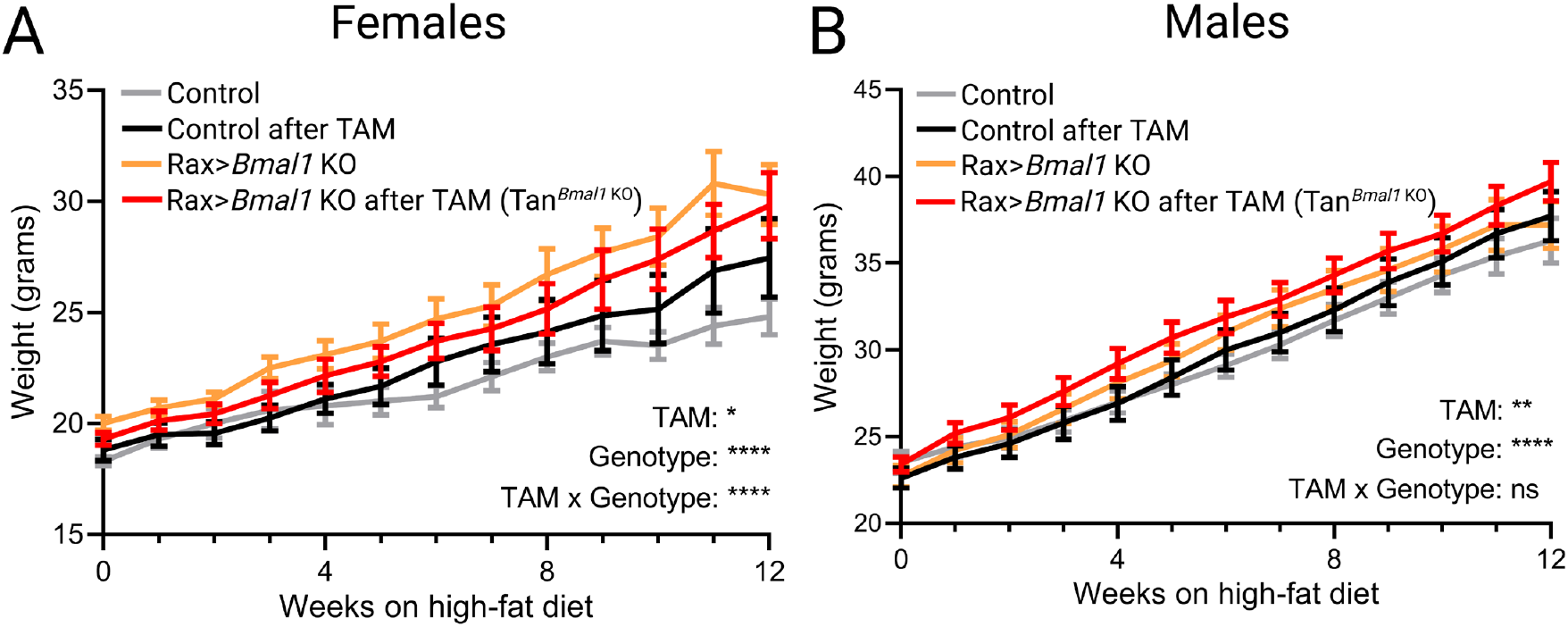
Adult tanycyte *Bmal1* KO reduces weight gain in female but not male mice on HFD. (**A-B)** Tan^*Bmal1* KO^ blocks weight gain (red vs orange) relative to genetic controls (black vs gray) on HFD in females (n = 10-15 mice/group), but not in males (n = 15-22 mice/group). Mean ± SEM, *p<0.05, **p<0.01, ****p<0.0001, ns = not significant, 3-way ANOVA.

In contrast, males exhibited main effects of both genotype (p<0.0001) and previous TAM/control treatment (p<0.01), but with no significant interaction. TAM treatment was associated with roughly similar weight gain in all genotypes in males, suggesting a drug effect independent of tanycyte Bmal1 status. In body composition analysis conducted after HFD feeding, both sexes had no significant effect on percent body weight derived from fat (Figure S2).

Importantly, given tanycytes’ known involvement in seasonal regulation of activity (Wood and Loudon, 2014), loss of tanycyte *Bmal1* did not alter the period or χ^2^ amplitude of wheel-running locomotor rhythms (Figure S3). Since disturbances in sleep are frequently associated with metabolic dysregulation (Depner et al., 2014; Scheer et al., 2009), we also tested for differences in sleep/wake behavior using the PiezoSleep system. Neither the circadian timing nor total amount of sleep/wake was affected by tanycyte-specific *Bmal1* knockout in either sex (Figure S4).

### Female mice have higher baseline adult tanycyte-derived ARC neurogenesis than males

From our sequencing analysis (Figure 1A-B), we suspected that tanycyte-derived ARC neurogenesis might contribute to attenuated weight gain on HFD in female Tan^*Bmal1* KO^ mice. But whether there is a baseline sex difference in tanycyte neurogenesis was unclear, with the two existing studies reporting no sex effect (Lee et al., 2014) and a weak trend toward increased female neurogenesis (Xu et al., 2005) in the hypothalamus. Moreover, both studies lack lineage tracing and are susceptible to uneven incorporation of BrdU by dividing cells, leaving ambiguous what fraction of this neurogenesis is truly tanycyte-derived.

To fill this gap, we administered TAM chow to female and male *RaxCreER*^*/+*^*;Ai14*^*/+*^ reporter mice, permanently labeling tanycytes and their progeny with lox-stopped tdTomato in adulthood (Madisen et al., 2010). ∼1-month after TAM induction, tdTomato-tagged ARC cells were then counted and categorized as neurons or astrocytes by morphology (Figure 2A). We observed higher baseline tanycyte-derived neurogenesis in female compared to male ARC using our system (Figure 3B-C). In contrast, tanycyte-derived ARC astrogenesis and neuron/astrocyte ratio had no sex difference.

**Figure 3.**
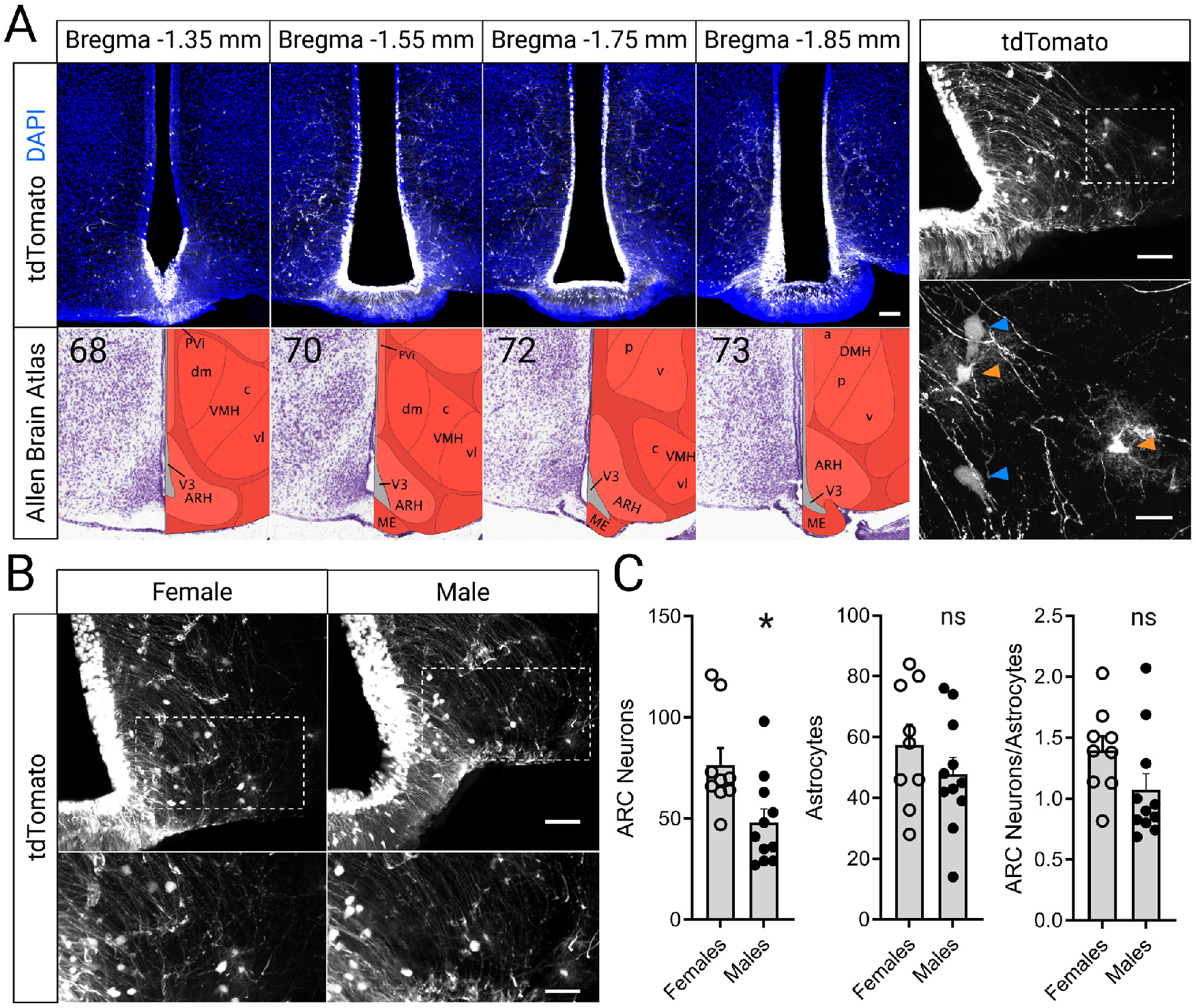
Lineage tracing reveals sexual dimorphism of tanycyte adult neurogenesis. (**A**) Left: representative 100μm hypothalamus slices used for lineage tracing cell counts (top), matched to corresponding Allen Brain Atlas images (bottom). Scale bar = 100μm. Right: representative slice (top) and magnified region of interest (bottom) showing tanycyte-derived ARC neurons (blue arrows) and astrocytes (orange arrows). Scale bars = 100, 75μm. (**B**) Representative coronal slices (top) and magnified regions of interest (bottom) of ARC neurons and astrocytes born from tanycytes in female and male mice. Scale bars = 100, 75μm. (**C**) Quantification of adult-born ARC neurons (left), astrocytes (middle), and neuron/astrocyte ratio derived from tanycytes in female and male mice (n = 9-11 mice/group). Mean + SEM. *p<0.05, ns = not significant, Student’s t-test.

### Tanycyte-derived adult ARC neurogenesis is sex-specifically reduced by adult *Bmal1* deletion in females but not males, reflecting depletion of adult-born orexinergic neurons

Having established this baseline, we next did similar lineage tracing in TAM-induced tanycyte *Bmal1*-deficient Tan^*Bmal1* KO^;*Ai14*^*/+*^ and genetic control *RaxCreER*^*/+*^*;Bmal1*^*lox/+*^*;Ai14*^*/+*^ ARC, in both females and males. Similarly to reduced weight gain we observed in these mice on HFD (Figure 2), adult tanycyte-derived neurogenesis was reduced in female but not male Tan^*Bmal1* KO^;*Ai14*^*/+*^ ARC (Figure 4A-D). In contrast, adult tanycyte-derived astrogenesis was increased in both female and male Tan^*Bmal1* KO^;*Ai14*^*/+*^ mice, leading to a significantly reduced adult tanycyte-derived neuron/astrocyte ratio only in females (Figure 4C-D).

**Figure 4.**
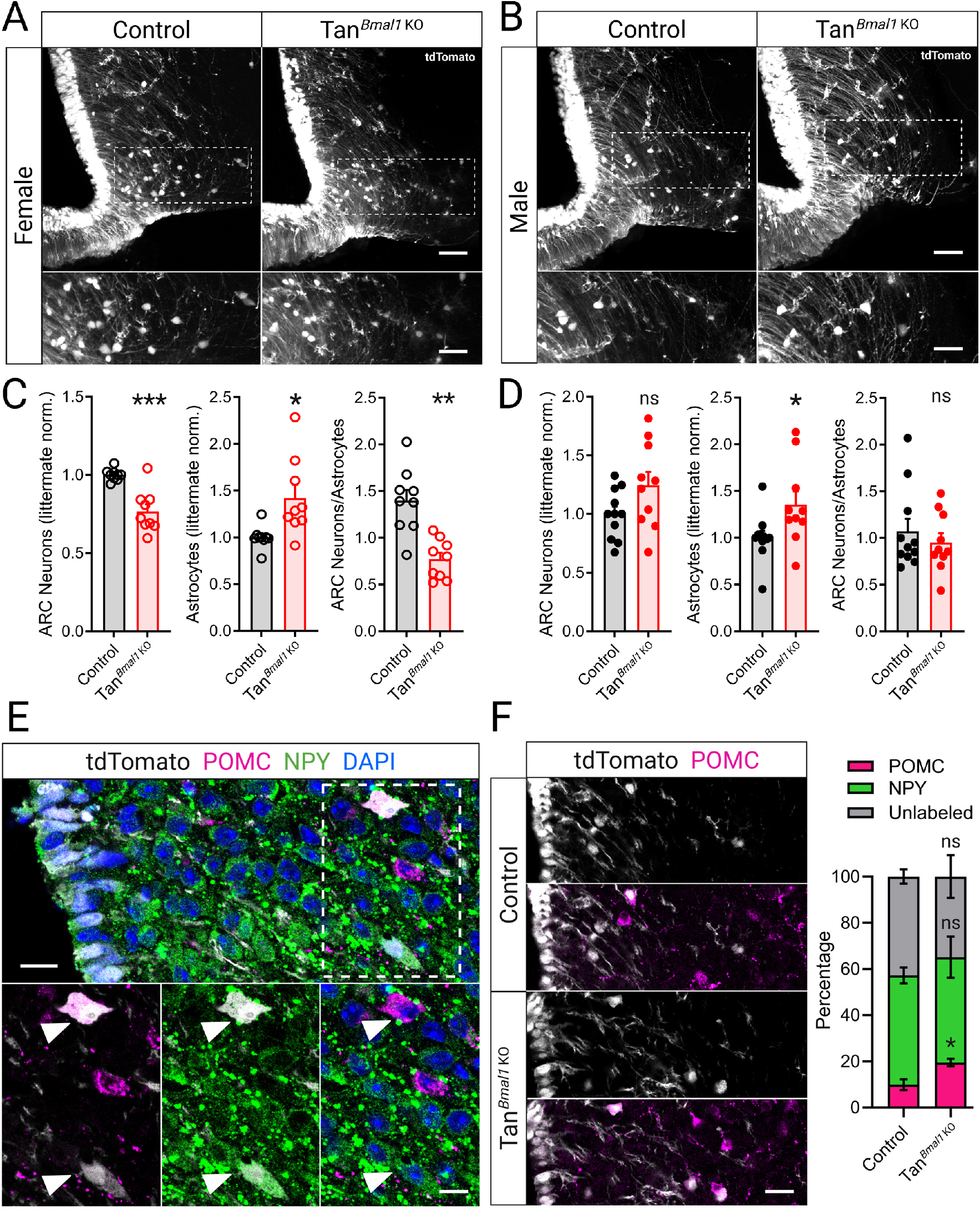
Impact of *Bmal1* KO on tanycyte neurogenesis is sexually dimorphic. (**A-B**) Representative coronal sections (top) and magnified regions of interest (bottom) showing adult-born ARC neurons and astrocytes derived from tanycytes in control and Tan^*Bmal1* KO^ mice. Scale bars = 100, 75μm. (**C-D**) Quantification of adult-born ARC neurons (left), astrocytes (middle), and neuron/astrocyte ratio (right) of tanycyte-derived cells from control and Tan^*Bmal1* KO^ mice (n = 9 female and 10-11 male mice/group). (**E**) Representative z-planes (top) and magnified regions of interest (bottom) with white arrows indicating POMC^+^ satiety (magenta) and NPY^+^ hunger (green) neurons lineage traced from tanycytes (white). Blue indicates Hoechst^+^ nuclei. Scale bars = 30μm. (**F**) Representative z-planes (left) and quantification (right) of tanycyte-derived ARC neuron fates in control and Tan^*Bmal1* KO^ mice (n=4-5 female mice/group). Scale bar = 50μm. Mean ± SEM, *p<0.05, **p<0.01, ***p<0.001, ns = not significant, *p<0.05, Student’s t-test.

The ARC contains both orexinergic AgRP/NPY^+^ and anorexinergic POMC^+^ neuropeptidergic neurons that are specified at a late clock-competent stage, and there is considerable evidence for a circadian clock role in specifying cell fate (Bedont et al., 2020, 2015). We hypothesized that reduced female-specific ARC neurogenesis caused by tanycyte *Bmal1* deletion (Figure 4) might reflect selective depletion of adult-born orexinergic ARC neurons, potentially contributing to reduced female weight gain on HFD (Figure 2). To investigate this possibility, we carried out lineage tracing in female mice with an extended 12-week chase (mirroring our HFD duration in Figure 2) and co-stained ARC for NPY, and POMC. While there was no effect of tanycyte *Bmal1* deletion on the percentage of adult-born, tanycyte-derived NPY^+^ or NPY/POMC-negative ARC neurons, the proportion of adult-born POMC^+^ neurons was increased (Figure 4E-F). Together with decreased overall tanycyte neurogenesis, this is consistent with a skewing of tanycyte-derived ARC neuronal fate toward anorexigenic fate after adult tanycyte *Bmal1* deletion.

## Discussion

In this work, we leveraged the RaxCreER driver to execute the first relatively specific *Bmal1* knockout in tanycytes *in vivo* and investigated the impact of this manipulation on diet-induced weight gain and hypothalamic adult neurogenesis. RaxCreER is well-characterized, and aside from hypothalamic tanycytes, targets only retinal Muller glia, cerebellum, and a handful of posterior pituicytes in adulthood (Pak et al., 2014). *Bmal1* deletion in these off-target cell-types is unlikely to drive the phenotypes we observe, not least because of attenuated HFD weight gain in female but not male mice after adult RaxCreER-mediated *Bmal1* deletion (Figure 2). This beautifully phenocopies female-specific attenuated HFD weight gain after focal irradiation-induced ablation of tanycyte neurogenic capacity (Lee et al., 2014, 2012). Thus, we interpret female-specific reduction of tanycyte-derived adult ARC neurogenesis after tanycyte *Bmal1* deletion (Figure 3) as a brake on female weight gain. Consistent with this, tanycyte-derived ARC neuron identities after tanycyte *Bmal1* deletion shift toward anorexigenic POMC^+^ fates (Figure 4). Notably, adult-born POMC^+^ neurons can partially rescue feeding and metabolism on a *Pomc*-deficient background (Surbhi et al., 2021), highlighting potential functional relevance of our anorexigenic shift.

This model is coherent with the circadian clock’s role in mature ARC circuitry. While HFD only modestly impairs ARC core clock gene circadian rhythmicity, it dramatically weakens circadian rhythmicity of genes controlled by ARC peptides, together with overall decreased orexigenic (*Npy/Agrp/Hypocretin*) and increased anorexigenic (*Pomc/Cart*) mRNA expression (Clemenzi et al., 2020; Kohsaka et al., 2007; Pendergast et al., 2013). Bmal1 is also required for acute palmitate induced *Npy* expression in a hypothalamic cell line (Clemenzi et al., 2020). In this context, skewing of adult-born neurons toward anorexigenic POMC^+^ fate after tanycyte *Bmal1* deletion (Figure 4) suggests a pro-orexigenic clock role across the developmental lifespan of female ARC neurons. Circadian Bmal1 competition for cis binding elements may contribute, as several other bHLH transcription factors promote anorexigenic fate during ARC development (Bedont et al., 2015). It is also possible that migration and/or survival effects contribute to circadian regulation of adult-born orexigenic/anorexigenic ratios.

Importantly, our findings for tanycyte-derived ARC neurogenesis are consistent with previously studied circadian clock roles elsewhere, predominantly the hippocampus. Hippocampal neural stem cells divide on a circadian rhythm, hyper-proliferate in clock-deficient young animals, and hypo-proliferate in clock-deficient old animals (Ali et al., 2015; Bouchard-Cannon et al., 2013; Encinas et al., 2011; Schnell et al., 2014). This likely reflects circadian clock roles in inhibiting excessive cell division and promoting self-renewal in young animals. In addition, Bmal1 has a potentially clock-independent role in promoting neural over glial daughter cell fate (Bedont et al., 2020). Accordingly, sex-agnostic increased ARC astrogenesis after tanycyte *Bmal1*-deletion suggests that Bmal1 specific pro-neural effects are sex independent (Figure 4). Conversely, female-specific decreased ARC neurogenesis from *Bmal1*-deficient tanycytes suggests that Bmal1’s role in restraining excessive cell division and promoting self-renewal is sexually dimorphic in tanycytes (Figure 4).

A hormonal interaction may contribute to sexually dimorphic tanycyte responses to *Bmal1* deletion. Indeed, estrogens require tanycyte ERα to exert anorexigenic effects (Fernandois et al., 2024), which include reduced weight gain on HFD and regulation of ARC neurogenesis and adult-born cell fate in female mice (Bless et al., 2016, 2014). The circadian clock may gate tanycyte sensitivity to estrogen signaling via Per2 regulation of ERα degradation (Gery et al., 2007), Clock interactions with ERα (Li et al., 2013), and/or other mechanisms.

Finally, while we focus on ARC neurogenesis as one likely mechanism, we do not discount the possibility that other factors may also contribute to regulation of female weight homeostasis by tanycyte Bmal1. Possibilities include modulation of tanycytic gating of hormone release, modification of brain hormones, barrier functions gating peripheral signals, and/or active reception or transport of peripheral signals (Friedman, 2019; Müller-Fielitz et al., 2017; Prevot et al., 2018; Rodriguez et al., 2005; Wood and Loudon, 2014).And while parallels to the hippocampal neurogenesis literature suggest that our female-specific tanycyte neurogenesis effects reflect disruption of the cell-autonomous circadian clock (see above), we cannot fully exclude possible non-circadian Bmal1 contributions to our female-specific tanycyte deletion phenotypes. This invites future exploration of the local clock’s role in tanycyte function, including well-documented roles in stress response, reproduction, photoperiodism, and more (Fernandois et al., 2024; Prevot et al., 2018; Wood and Loudon, 2014). Indeed, even behavioral outputs such as circadian locomotor rhythms and sleep behavior that were unaffected in our study (Figures S3-4) may well be regulated by the tanycyte clock in the context of seasonality. Looking forward, we believe that the tanycyte clock’s impacts on physiology and behavior are a largely untapped field ripe for future study.

## Acknowledgments

We thank Dr. Seth Blackshaw for providing the *RaxCreER* mice, and helpful comments on the manuscript. We thank Dr. John Campbell for generously sharing an annotated Seurat gene expression library from his previous work (Campbell et al., 2017). We thank Dr. W. Timothy O’Brien and the Neurobehavior Testing Core at UPenn/ITMAT and IDDRC at CHOP/Penn U54 HD086984 for assistance with behavior procedures. We thank Dr. Corey Holman and the Rodent Metabolic Phenotyping Core (RRID: SCR_022427), supported in part by NIH grant S10-OD025098, the Cox Institute, and the Institute for Diabetes, Obesity and Metabolism at the University of Pennsylvania, for performing body composition analysis. This work was supported by grants from the NIH: F32MH125600 (to D.M.I.), 5R01AG068577 (to R.C.A.), F32AG056081, K99/R00NS118561 (to J.L.B), R37NS048471 (to A.S.), and the Howard Hughes Medical Institute (to A.S.). Figures were generated using BioRender. This article is subject to HHMI’s Open Access to Publications policy. HHMI lab heads have previously granted a nonexclusive CC BY 4.0 license to the public and a sublicensable license to HHMI in their research articles. Pursuant to those licenses, the author-accepted manuscript of this article can be made freely available under a CC BY 4.0 license immediately upon publication.

## Author contributions

D.M.I., J.L.B., and A.S. designed research; D.M.I., P.P., M.L.V., S.B.N., H.L., and J.L.B. performed research; J.Y. performed RNA-seq analyses with guidance from R.C.A.; D.M.I., M.L.V., and J.L.B. analyzed data; D.M.I. and J.L.B. wrote the paper; all authors intellectually contributed to the paper.

## Bibliography

Ali, A.A.H., Schwarz-Herzke, B., Stahr, A., Prozorovski, T., Aktas, O., Von Gall, C., 2015. Premature aging of the hippocampal neurogenic niche in adult Bmal1-deficient mice. Aging 7, 435–449. 10.18632/aging.100764

Astiz, M., Heyde, I., Oster, H., 2019. Mechanisms of Communication in the Mammalian Circadian Timing System. IJMS 20, 343. 10.3390/ijms20020343

Balland, E., Dam, J., Langlet, F., Caron, E., Steculorum, S., Messina, A., Rasika, S., Falluel-Morel, A., Anouar, Y., Dehouck, B., Trinquet, E., Jockers, R., Bouret, S.G., Prévot, V., 2014. Hypothalamic Tanycytes Are an ERK-Gated Conduit for Leptin into the Brain. Cell Metabolism 19, 293–301. 10.1016/j.cmet.2013.12.015

Bedont, J.L., Iascone, D.M., Sehgal, A., 2020. The Lineage Before Time: Circadian and Nonclassical Clock Influences on Development. Annu. Rev. Cell Dev. Biol. 36, 469–509. 10.1146/annurev-cellbio-100818-125454

Bedont, J.L., LeGates, T.A., Slat, E.A., Byerly, M.S., Wang, H., Hu, J., Rupp, A.C., Qian, J., Wong, G.W., Herzog, E.D., Hattar, S., Blackshaw, S., 2014. Lhx1 Controls Terminal Differentiation and Circadian Function of the Suprachiasmatic Nucleus. Cell Reports 7, 609–622. 10.1016/j.celrep.2014.03.060

Bedont, J.L., Newman, E.A., Blackshaw, S., 2015. Patterning, specification, and differentiation in the developing hypothalamus. WIREs Developmental Biology 4, 445–468. 10.1002/wdev.187

Bedont, J.L., Rohr, K.E., Bathini, A., Hattar, S., Blackshaw, S., Sehgal, A., Evans, J.A., 2018. Asymmetric vasopressin signaling spatially organizes the master circadian clock. J of Comparative Neurology 526, 2048–2067. 10.1002/cne.24478

Bless, E.P., Reddy, T., Acharya, K.D., Beltz, B.S., Tetel, M.J., 2014. Oestradiol and Diet Modulate Energy Homeostasis and Hypothalamic Neurogenesis in the Adult Female Mouse. J Neuroendocrinology 26, 805–816. 10.1111/jne.12206

Bless, E.P., Yang, J., Acharya, K.D., Nettles, S.A., Vassoler, F.M., Byrnes, E.M., Tetel, M.J., 2016. Adult Neurogenesis in the Female Mouse Hypothalamus: Estradiol and High-Fat Diet Alter the Generation of Newborn Neurons Expressing Estrogen Receptor α. eneuro 3, ENEURO.0027-16.2016. 10.1523/ENEURO.0027-16.2016

Bouchard-Cannon, P., Mendoza-Viveros, L., Yuen, A., Kærn, M., Cheng, H.-Y.M., 2013. The Circadian Molecular Clock Regulates Adult Hippocampal Neurogenesis by Controlling the Timing of Cell-Cycle Entry and Exit. Cell Reports 5, 961–973. 10.1016/j.celrep.2013.10.037

Campbell, J.N., Macosko, E.Z., Fenselau, H., Pers, T.H., Lyubetskaya, A., Tenen, D., Goldman, M., Verstegen, A.M.J., Resch, J.M., McCarroll, S.A., Rosen, E.D., Lowell, B.B., Tsai, L.T., 2017. A molecular census of arcuate hypothalamus and median eminence cell types. Nat Neurosci 20, 484–496. 10.1038/nn.4495

Chaker, Z., George, C., Petrovska, M., Caron, J.-B., Lacube, P., Caillé, I., Holzenberger, M., 2016. Hypothalamic neurogenesis persists in the aging brain and is controlled by energy-sensing IGF-I pathway. Neurobiology of Aging 41, 64–72. 10.1016/j.neurobiolaging.2016.02.008

Clemenzi, M.N., Martchenko, A., Loganathan, N., Tse, E.K., Brubaker, P.L., Belsham, D.D., 2020. Analysis of Western diet, palmitate and BMAL1 regulation of neuropeptide Y expression in the murine hypothalamus and BMAL1 knockout cell models. Molecular and Cellular Endocrinology 507, 110773. 10.1016/j.mce.2020.110773

Correa-da-Silva, F., Fliers, E., Swaab, D.F., Yi, C., 2021. Hypothalamic neuropeptides and neurocircuitries in Prader Willi syndrome. J Neuroendocrinology 33, e12994. 10.1111/jne.12994

Depner, C.M., Stothard, E.R., Wright, K.P., 2014. Metabolic Consequences of Sleep and Circadian Disorders. Curr Diab Rep 14, 507. 10.1007/s11892-014-0507-z

Encinas, J.M., Michurina, T.V., Peunova, N., Park, J.-H., Tordo, J., Peterson, D.A., Fishell, G., Koulakov, A., Enikolopov, G., 2011. Division-Coupled Astrocytic Differentiation and Age-Related Depletion of Neural Stem Cells in the Adult Hippocampus. Cell Stem Cell 8, 566–579. 10.1016/j.stem.2011.03.010

Fernandois, D., Rusidzé, M., Mueller-Fielitz, H., Sauve, F., Deligia, E., Silva, M.S.B., Evrard, F., Franco-García, A., Mazur, D., Martinez-Corral, I., Jouy, N., Rasika, S., Maurage, C.-A., Giacobini, P., Nogueiras, R., Dehouck, B., Schwaninger, M., Lenfant, F., Prevot, V., 2024. Estrogen receptor-α signaling in tanycytes lies at the crossroads of fertility and metabolism. Metabolism 158, 155976. 10.1016/j.metabol.2024.155976

Flores, A.E., Flores, J.E., Deshpande, H., Picazo, J.A., Xinmin Xie, Franken, P., Heller, H.C., Grahn, D.A., O’Hara, B.F., 2007. Pattern Recognition of Sleep in Rodents Using Piezoelectric Signals Generated by Gross Body Movements. IEEE Trans. Biomed. Eng. 54, 225–233. 10.1109/TBME.2006.886938

Frank, C.L., Ge, X., Xie, Z., Zhou, Y., Tsai, L.-H., 2010. Control of Activating Transcription Factor 4 (ATF4) Persistence by Multisite Phosphorylation Impacts Cell Cycle Progression and Neurogenesis*. Journal of Biological Chemistry 285, 33324–33337. 10.1074/jbc.M110.140699

Friedman, J.M., 2019. Leptin and the endocrine control of energy balance. Nat Metab 1, 754–764. 10.1038/s42255-019-0095-y

Fusco, C.M., Desch, K., Dörrbaum, A.R., Wang, M., Staab, A., Chan, I.C.W., Vail, E., Villeri, V., Langer, J.D., Schuman, E.M., 2021. Neuronal ribosomes exhibit dynamic and context-dependent exchange of ribosomal proteins. Nat Commun 12, 6127. 10.1038/s41467-021-26365-x

Gery, S., Virk, R.K., Chumakov, K., Yu, A., Koeffler, H.P., 2007. The clock gene Per2 links the circadian system to the estrogen receptor. Oncogene 26, 7916–7920. 10.1038/sj.onc.1210585

Guilding, C., Hughes, A.T., Brown, T.M., Namvar, S., Piggins, H.D., 2009. A riot of rhythms: neuronal and glial circadian oscillators in the mediobasal hypothalamus. Mol Brain 2, 28. 10.1186/1756-6606-2-28

Guo, H., Brewer, J.M., Champhekar, A., Harris, R.B.S., Bittman, E.L., 2005. Differential control of peripheral circadian rhythms by suprachiasmatic-dependent neural signals. Proc. Natl. Acad. Sci. U.S.A. 102, 3111–3116. 10.1073/pnas.0409734102

Haan, N., Goodman, T., Najdi-Samiei, A., Stratford, C.M., Rice, R., El Agha, E., Bellusci, S., Hajihosseini, M.K., 2013. Fgf10-Expressing Tanycytes Add New Neurons to the Appetite/Energy-Balance Regulating Centers of the Postnatal and Adult Hypothalamus. J. Neurosci. 33, 6170–6180. 10.1523/JNEUROSCI.2437-12.2013

Hastings, M.H., Maywood, E.S., Brancaccio, M., 2018. Generation of circadian rhythms in the suprachiasmatic nucleus. Nat Rev Neurosci 19, 453–469. 10.1038/s41583-018-0026-z

Herichová, I., Hasáková, K., Lukáčová, D., Mravec, B., Horváthová, Ľ., Kavická, D., 2017. Prefrontal Cortex and Dorsomedial Hypothalamus Mediate Food Reward-Induced Effects via npas2 and egr1 Expression in Rat. Physiol Res S501–S510. 10.33549/physiolres.933799

Hughes, M.E., Hogenesch, J.B., Kornacker, K., 2010. JTK_CYCLE: An Efficient Nonparametric Algorithm for Detecting Rhythmic Components in Genome-Scale Data Sets. J Biol Rhythms 25, 372–380. 10.1177/0748730410379711

Kawashima, F., Saito, K., Kurata, H., Maegaki, Y., Mori, T., 2017. c-jun is differentially expressed in embryonic and adult neural precursor cells. Histochem Cell Biol 147, 721–731. 10.1007/s00418-016-1536-2

Kohsaka, A., Laposky, A.D., Ramsey, K.M., Estrada, C., Joshu, C., Kobayashi, Y., Turek, F.W., Bass, J., 2007. High-Fat Diet Disrupts Behavioral and Molecular Circadian Rhythms in Mice. Cell Metabolism 6, 414– 421. 10.1016/j.cmet.2007.09.006

Lee, D.A., Bedont, J.L., Pak, T., Wang, H., Song, J., Miranda-Angulo, A., Takiar, V., Charubhumi, V., Balordi, F., Takebayashi, H., Aja, S., Ford, E., Fishell, G., Blackshaw, S., 2012. Tanycytes of the hypothalamic median eminence form a diet-responsive neurogenic niche. Nat Neurosci 15, 700–702. 10.1038/nn.3079

Lee, D.A., Yoo, S., Pak, T., Salvatierra, J., Velarde, E., Aja, S., Blackshaw, S., 2014. Dietary and sex-specific factors regulate hypothalamic neurogenesis in young adult mice. Front. Neurosci. 8. 10.3389/fnins.2014.00157

Lein, E.S., Hawrylycz, M.J., Ao, N., Ayres, M., Bensinger, A., Bernard, A., Boe, A.F., Boguski, M.S., Brockway, K.S., Byrnes, E.J., Chen, Lin, Chen, Li, Chen, T.-M., Chi Chin, M., Chong, J., Crook, B.E., Czaplinska, A., Dang, C.N., Datta, S., Dee, N.R., Desaki, A.L., Desta, T., Diep, E., Dolbeare, T.A., Donelan, M.J., Dong, H.-W., Dougherty, J.G., Duncan, B.J., Ebbert, A.J., Eichele, G., Estin, L.K., Faber, C., Facer, B.A., Fields, R., Fischer, S.R., Fliss, T.P., Frensley, C., Gates, S.N., Glattfelder, K.J., Halverson, K.R., Hart, M.R., Hohmann, J.G., Howell, M.P., Jeung, D.P., Johnson, R.A., Karr, P.T., Kawal, R., Kidney, J.M., Knapik, R.H., Kuan, C.L., Lake, J.H., Laramee, A.R., Larsen, K.D., Lau, C., Lemon, T.A., Liang, A.J., Liu, Y., Luong, L.T., Michaels, J., Morgan, J.J., Morgan, R.J., Mortrud, M.T., Mosqueda, N.F., Ng, L.L., Ng, R., Orta, G.J., Overly, C.C., Pak, T.H., Parry, S.E., Pathak, S.D., Pearson, O.C., Puchalski, R.B., Riley, Z.L., Rockett, H.R., Rowland, S.A., Royall, J.J., Ruiz, M.J., Sarno, N.R., Schaffnit, K., Shapovalova, N.V., Sivisay, T., Slaughterbeck, C.R., Smith, S.C., Smith, K.A., Smith, B.I., Sodt, A.J., Stewart, N.N., Stumpf, K.-R., Sunkin, S.M., Sutram, M., Tam, A., Teemer, C.D., Thaller, C., Thompson, C.L., Varnam, L.R., Visel, A., Whitlock, R.M., Wohnoutka, P.E., Wolkey, C.K., Wong, V.Y., Wood, M., Yaylaoglu, M.B., Young, R.C., Youngstrom, B.L., Feng Yuan, X., Zhang, B., Zwingman, T.A., Jones, A.R., 2007. Genome-wide atlas of gene expression in the adult mouse brain. Nature 445, 168–176. 10.1038/nature05453

Li, S., Wang, M., Ao, X., Chang, A.K., Yang, C., Zhao, F., Bi, H., Liu, Y., Xiao, L., Wu, H., 2013. CLOCK is a substrate of SUMO and sumoylation of CLOCK upregulates the transcriptional activity of estrogen receptor-α. Oncogene 32, 4883–4891. 10.1038/onc.2012.518

Lu, R., Dong, Y., Li, J.-D., 2020. Necdin regulates BMAL1 stability and circadian clock through SGT1-HSP90 chaperone machinery. Nucleic Acids Research 48, 7944–7957. 10.1093/nar/gkaa601

Ma, Y., Cheng, Q., Ren, Z., Xu, L., Zhao, Y., Sun, J., Hu, S., Xiao, W., 2012. Induction of IGF-1R expression by EGR-1 facilitates the growth of prostate cancer cells. Cancer Letters 317, 150–156. 10.1016/j.canlet.2011.11.021

Madisen, L., Zwingman, T.A., Sunkin, S.M., Oh, S.W., Zariwala, H.A., Gu, H., Ng, L.L., Palmiter, R.D., Hawrylycz, M.J., Jones, A.R., Lein, E.S., Zeng, H., 2010. A robust and high-throughput Cre reporting and characterization system for the whole mouse brain. Nat Neurosci 13, 133–140. 10.1038/nn.2467

Mang, G.M., Nicod, J., Emmenegger, Y., Donohue, K.D., O’Hara, B.F., Franken, P., 2014. Evaluation of a Piezoelectric System as an Alternative to Electroencephalogram/ Electromyogram Recordings in Mouse Sleep Studies. Sleep 37, 1383–1392. 10.5665/sleep.3936

Martínez-Zamudio, R.I., Roux, P.-F., De Freitas, J.A.N.L.F., Robinson, L., Doré, G., Sun, B., Belenki, D., Milanovic, M., Herbig, U., Schmitt, C.A., Gil, J., Bischof, O., 2020. AP-1 imprints a reversible transcriptional programme of senescent cells. Nat Cell Biol 22, 842–855. 10.1038/s41556-020-0529-5

McMenamin, T., 2007. A time to work: recent trends in shift work and flexible schedules. Bureau of Labor Statistics.

Moffat, J.J., Ka, M., Jung, E.-M., Smith, A.L., Kim, W.-Y., 2017. The role of MACF1 in nervous system development and maintenance. Seminars in Cell & Developmental Biology 69, 9–17. 10.1016/j.semcdb.2017.05.020

Mohawk, J.A., Green, C.B., Takahashi, J.S., 2012. Central and Peripheral Circadian Clocks in Mammals. Annu. Rev. Neurosci. 35, 445–462. 10.1146/annurev-neuro-060909-153128

Müller-Fielitz, H., Stahr, M., Bernau, M., Richter, M., Abele, S., Krajka, V., Benzin, A., Wenzel, J., Kalies, K., Mittag, J., Heuer, H., Offermanns, S., Schwaninger, M., 2017. Tanycytes control the hormonal output of the hypothalamic-pituitary-thyroid axis. Nat Commun 8, 484. 10.1038/s41467-017-00604-6

Nordhaus, W., 1996. Do Real-Output and Real-Wage Measures Capture Reality? The History of Lighting Suggests Not, in: The Economics of New Goods. National Bureau of Economic Research, Inc., pp. 27– 70.

Pak, T., Yoo, S., Miranda-Angulo, A.M., Wang, H., Blackshaw, S., 2014. Rax-CreERT2 Knock-In Mice: A Tool for Selective and Conditional Gene Deletion in Progenitor Cells and Radial Glia of the Retina and Hypothalamus. PLoS ONE 9, e90381. 10.1371/journal.pone.0090381

Pakos-Zebrucka, K., Koryga, I., Mnich, K., Ljujic, M., Samali, A., Gorman, A.M., 2016. The integrated stress response. EMBO Reports 17, 1374–1395. 10.15252/embr.201642195

Pendergast, J.S., Branecky, K.L., Yang, W., Ellacott, K.L.J., Niswender, K.D., Yamazaki, S., 2013. High-fat diet acutely affects circadian organisation and eating behavior. Eur J of Neuroscience 37, 1350–1356. 10.1111/ejn.12133

Prevot, V., Dehouck, B., Sharif, A., Ciofi, P., Giacobini, P., Clasadonte, J., 2018. The Versatile Tanycyte: A Hypothalamic Integrator of Reproduction and Energy Metabolism. Endocrine Reviews 39, 333–368. 10.1210/er.2017-00235

Rodriguez, E., Blazquez, J., Pastor, F., Pelaez, B., Pena, P., Peruzzo, B., Amat, P., 2005. Hypothalamic Tanycytes: A Key Component of Brain–Endocrine Interaction. International Review of Cytology 247, 89–164. 10.1016/S0074-7696(05)47003-5

Rodríguez-Cortés, B., Hurtado-Alvarado, G., Martínez-Gómez, R., León-Mercado, L.A., Prager-Khoutorsky, M., Buijs, R.M., 2022. Suprachiasmatic nucleus-mediated glucose entry into the arcuate nucleus determines the daily rhythm in blood glycemia. Current Biology 32, 796-805.e4. 10.1016/j.cub.2021.12.039

Scheer, F.A.J.L., Hilton, M.F., Mantzoros, C.S., Shea, S.A., 2009. Adverse metabolic and cardiovascular consequences of circadian misalignment. Proc. Natl. Acad. Sci. U.S.A. 106, 4453–4458. 10.1073/pnas.0808180106

Schnell, A., Chappuis, S., Schmutz, I., Brai, E., Ripperger, J.A., Schaad, O., Welzl, H., Descombes, P., Alberi, L., Albrecht, U., 2014. The Nuclear Receptor REV-ERBα Regulates Fabp7 and Modulates Adult Hippocampal Neurogenesis. PLoS ONE 9, e99883. 10.1371/journal.pone.0099883

Shimogori, T., Lee, D.A., Miranda-Angulo, A., Yang, Y., Wang, H., Jiang, L., Yoshida, A.C., Kataoka, A., Mashiko, H., Avetisyan, M., Qi, L., Qian, J., Blackshaw, S., 2010. A genomic atlas of mouse hypothalamic development. Nat Neurosci 13, 767–775. 10.1038/nn.2545

Surbhi Wittmann, G., Low, M.J., Lechan, R.M., 2021. Adult-born proopiomelanocortin neurons derived from Rax-expressing precursors mitigate the metabolic effects of congenital hypothalamic proopiomelanocortin deficiency. Molecular Metabolism 53, 101312. 10.1016/j.molmet.2021.101312

Török, I., Herrmann-Horle, D., Kiss, I., Tick, G., Speer, G., Schmitt, R., Mechler, B.M., 1999. Down-Regulation of RpS21, a Putative Translation Initiation Factor Interacting with P40, Produces Viable Minute Imagos and Larval Lethality with Overgrown Hematopoietic Organs and Imaginal Discs. Molecular and Cellular Biology 19, 2308–2321. 10.1128/MCB.19.3.2308

Wood, S., Loudon, A., 2014. Clocks for all seasons: unwinding the roles and mechanisms of circadian and interval timers in the hypothalamus and pituitary. Journal of Endocrinology 222, R39–R59. 10.1530/JOE-14-0141

Wu, G., Anafi, R.C., Hughes, M.E., Kornacker, K., Hogenesch, J.B., 2016. MetaCycle: an integrated R package to evaluate periodicity in large scale data. Bioinformatics 32, 3351–3353. 10.1093/bioinformatics/btw405

Xu, Y., Tamamaki, N., Noda, T., Kimura, K., Itokazu, Y., Matsumoto, N., Dezawa, M., Ide, C., 2005. Neurogenesis in the ependymal layer of the adult rat 3rd ventricle. Experimental Neurology 192, 251– 264. 10.1016/j.expneurol.2004.12.021

Yasuo, S., Von Gall, C., Weaver, D.R., Korf, H., 2008. Rhythmic expression of clock genes in the ependymal cell layer of the third ventricle of rodents is independent of melatonin signaling. Eur J of Neuroscience 28, 2443–2450. 10.1111/j.1460-9568.2008.06541.x

Yi, J.S., Díaz, N.M., D’Souza, S., Buhr, E.D., 2022. The molecular clockwork of mammalian cells. Seminars in Cell & Developmental Biology 126, 87–96. 10.1016/j.semcdb.2021.03.012

Yoo, S., Blackshaw, S., 2018. Regulation and function of neurogenesis in the adult mammalian hypothalamus. Progress in Neurobiology 170, 53–66. 10.1016/j.pneurobio.2018.04.001

Yoo, S., Cha, D., Kim, S., Jiang, L., Cooke, P., Adebesin, M., Wolfe, A., Riddle, R., Aja, S., Blackshaw, S., 2020. Tanycyte ablation in the arcuate nucleus and median eminence increases obesity susceptibility by increasing body fat content in male mice. Glia 68, 1987–2000. 10.1002/glia.23817

Yoshikawa, K., 2021. Necdin: A purposive integrator of molecular interaction networks for mammalian neuron vitality. Genes to Cells 26, 641–683. 10.1111/gtc.12884

