## Supplemental Figures for "Tanycyte Bmal1 sex-specifically regulates weight gain and hypothalamic neurogenesis in female mice"

### Supplementary Figure 1

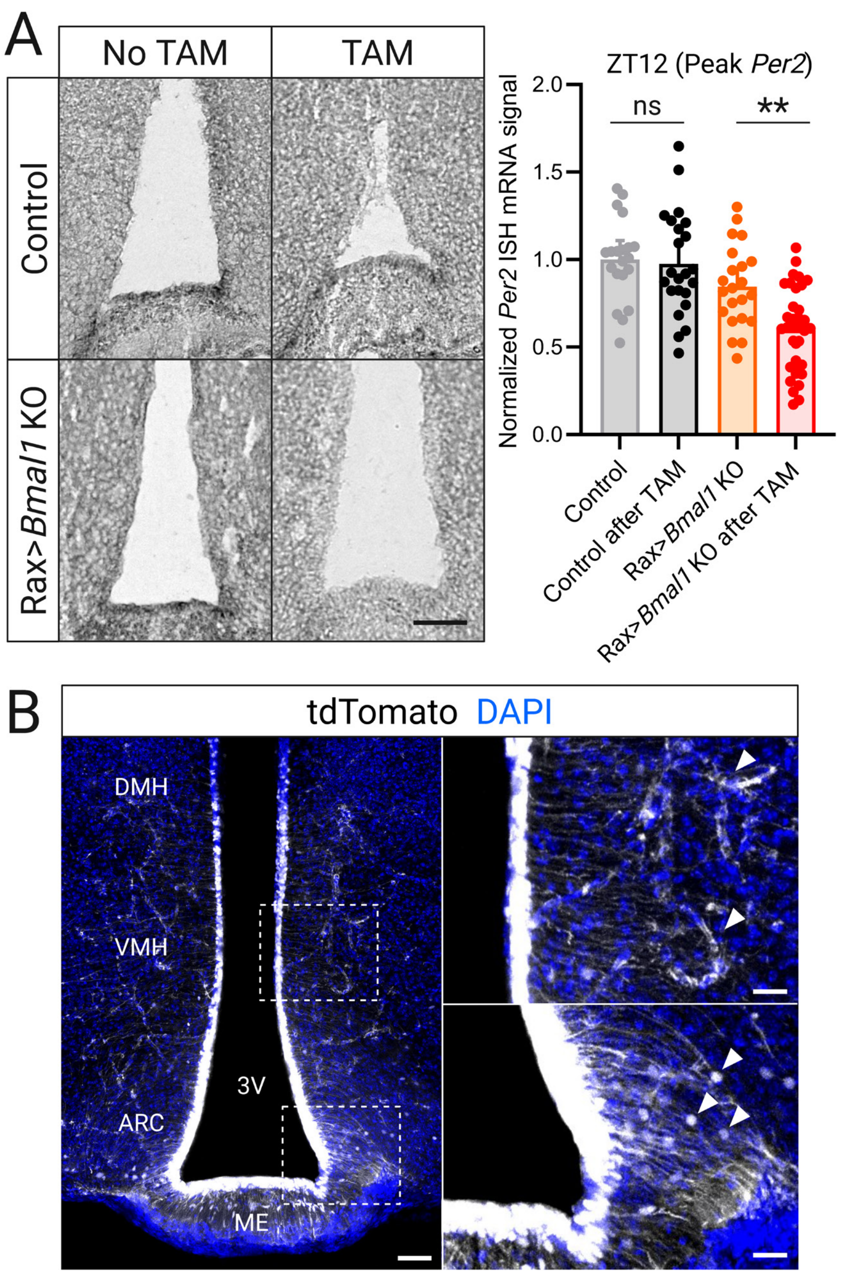


**Supplementary Figure 1. Tanycyte-specific *Bmal1* KO and lineage tracing with *RaxCreER* driver mice**
(**A**) Representative images (left) and quantification (right) of *Per2* mRNA levels at ZT12 in mice following 12 weeks of HFD demonstrates that tanycyte-specific *Bmal1* KO through tamoxifen (TAM)-induced recombination in *RaxCreER* mice reduces peak *Per2* expression (same mice as in Figure 3; n=20-35 mixed-sex mice per group). Scale bar = 150 µm. (**B**) Coronal brain section of mouse hypothalamus 35 days following RaxCreER-driven tdTomato expression (white; same image as Figure 3A panel 3 with magnified regions of interest for detailed examination). Dashed boxes indicate magnified regions with tanycyte processes contacting blood vessels (white arrows in top region of interest) and tanycyte-born neurons (white arrows in bottom region of interest). Blue indicates Hoechst+ nuclei. 3V third ventricle, ARC arcuate nucleus, DMH dorsomedial nucleus, ME median eminence, VMH ventromedial nucleus. Scale bars = 50µm (left), 20µm (right). Mean + SEM, **p<0.01, ns = not significant, two-way ANOVA with Holm–Sidak post hoc test.

Supplementary Figure 2

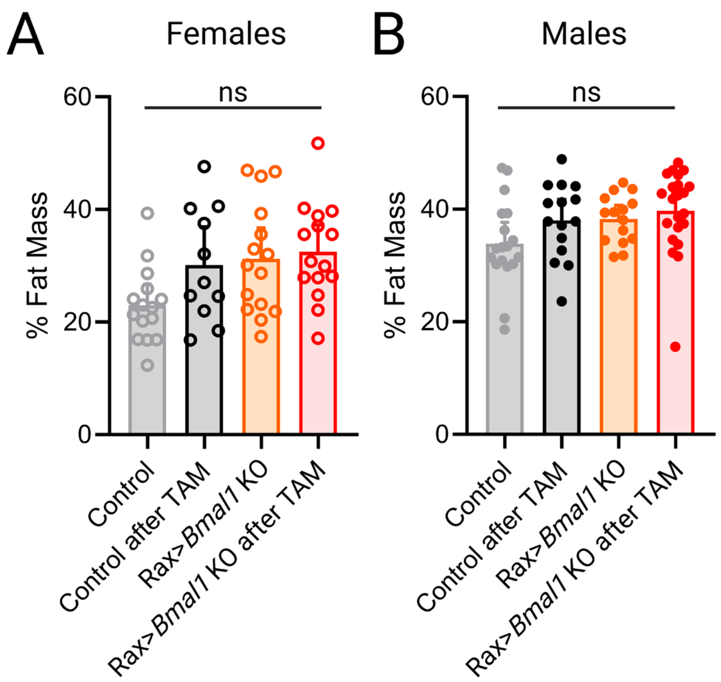


**Supplementary Figure 2. Tanycyte *Bmal1* KO does not alter fat mass composition following high-fat diet** (**A-B**) Percent fat mass measured with MRI of female (A) and male (B) mice following 12 weeks of HFD (n = 11-15 female and 15-22 male mice per group). Error bars indicate mean + SEM. ns = not significant, two-way ANOVA with Holm–Sidak post hoc test.

Supplementary Figure 3

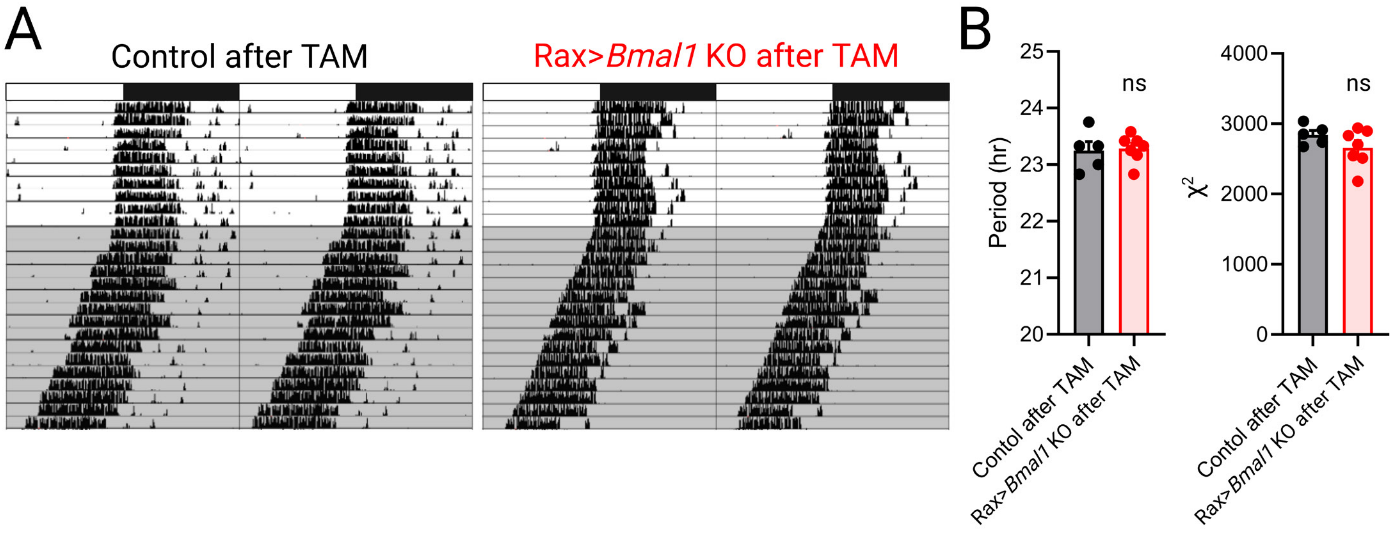


**Supplementary Figure 3. Tanycyte *Bmal1* KO does not alter circadian locomotor activity**
(**A**) Representative double-plotted running wheel actograms for control and Tan$^{Bmal1\ KO}$ mice after TAM. White and black bars denote 12:12-hour LD cycle for the first 10 days and the gray shaded area denotes DD (last 15 days). (**B**) Average circadian period (left) and $\chi^2$ amplitude (right) of wheel-running activity during 15 days of constant darkness (n = 5-7 mixed-sex mice per group). Mean + SEM. ns = not significant, Student's t-test.

### Supplementary Figure 4

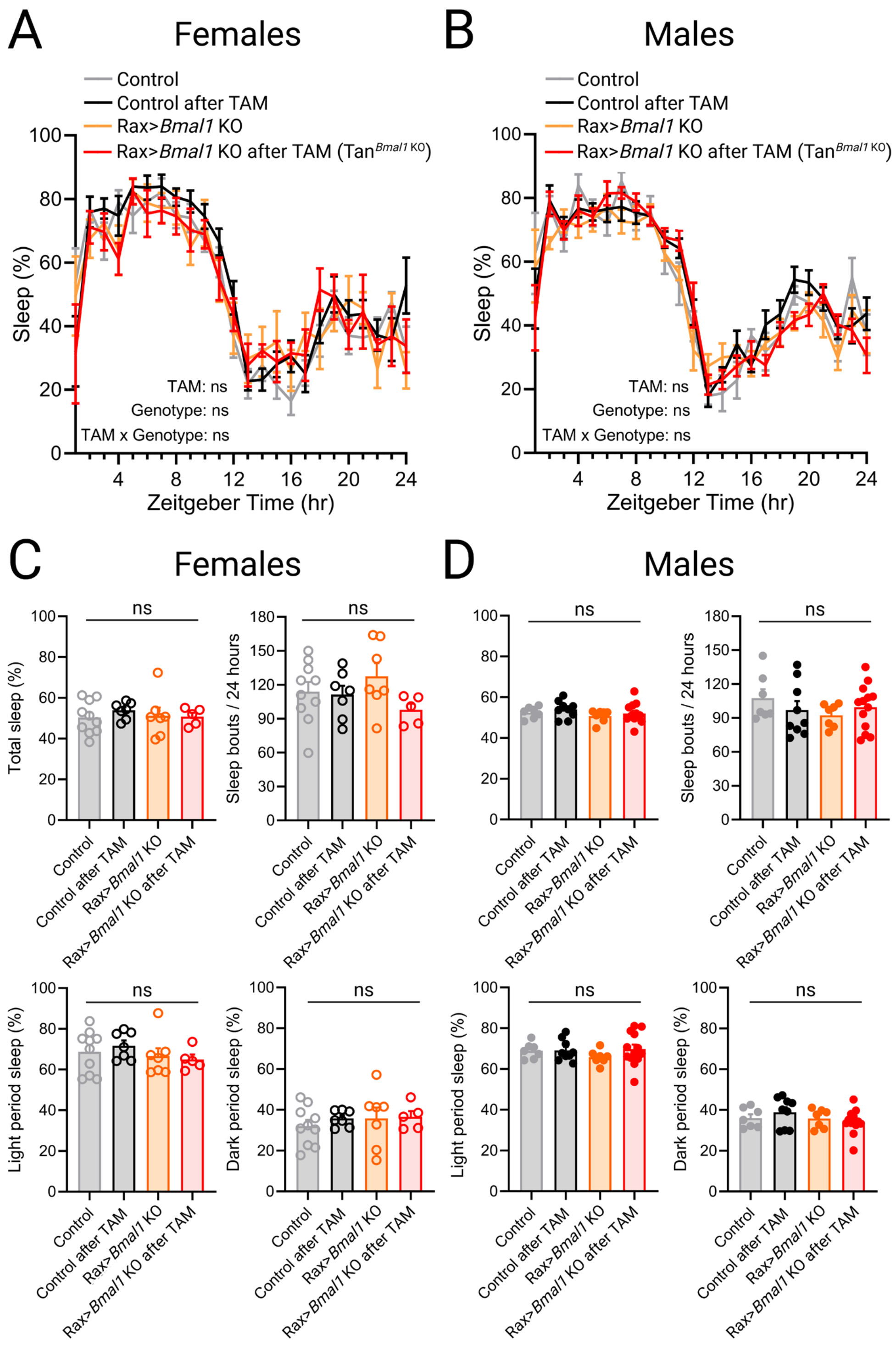


**Supplementary Figure 4. Tanycyte *Bmal1* KO does not alter sleep**
(**A-B**) Percent time asleep in female (A) and male (B) mice binned in 1-hour intervals averaged across 4 days (n = 5-10 female and 7-13 male mice/group). (**C-D**) Percent time asleep and sleep bouts/day (top) and percent time spent asleep during the light period and dark period (bottom) of female (C) and male (D) mice quantified from (A-B). Mean ± SEM. ns = not significant, three-way ANOVA (A-B) and two-way ANOVA with Holm–Sidak post hoc test (C-D).
